# Compensatory placental hypervascularization in a rat model of early-stage, diet-induced maternal obesity

**DOI:** 10.1101/2025.07.15.664837

**Authors:** Yhoiss Smiht Munoz Ceron, Sirsa Aleida Hidalgo Ibarra, David Moreno Martinez, Maria Eleonora Tejada Lopez, Jenniffer Alejandra Castellanos-Garzon, Liliana Salazar Monsalve, Maria Carolina Pustovrh

## Abstract

**Objective:** Maternal obesity is usually associated with placental hypovascularity. This study aimed to challenge this paradigm by investigating the immediate vascular adaptations in the placental labyrinth zone in response to short-term, diet-induced obesity. The study hypothesised that there would be an initial compensatory hypervascularisation before the onset of systemic metabolic disease.

**Methods:** Female Wistar rats were fed either a standard control diet (CG, n = 9) or an ultra-processed, hypercaloric cafeteria diet (EG, n = 9) for eight weeks to induce obesity. On gestational day 16.5, maternal morphometric and biochemical analyses were performed alongside detailed placental histomorphometry and immunohistochemistry for CD31/α-actin, in order to quantify foetal vessel density in the labyrinth zone using ImageJ.

**Results:** The cafeteria diet successfully induced a significant obese phenotype (mean weight gain: 62.73 g in the experimental group (EG) versus 32.26 g in the control group (CG); P < 0.0001), but did not induce significant hyperglycaemia or dyslipidaemia (P > 0.05). Although there were no significant differences in foetal or placental weights, the labyrinth zone of the EG showed a significant increase in foetal vessel density (29.08 ± 1.91 vessels/field) compared to the CG (26.06 ± 1.80 vessels/field; P = 0.014), indicating robust vascular remodelling.

**Conclusion:** Short-term exposure to an ‘obesogenic’ diet triggers significant compensatory hypervascularisation in the placenta of rats, which is an adaptive response that precedes systemic metabolic dysfunction. This finding contrasts with the hypovascularity observed in chronic obesity. The discrepancy between increased vascular density and foetal growth underscores the importance of evaluating not only the quantity but also the quality and function of blood vessels when assessing placental health in cases of maternal obesity.

**Highlights:**

- Acute cafeteria diet-induced obesity causes placental hypervascularity.
- Increased placental vasculature does not correlate with enhanced foetal growth.
- Rapid obesity develops without major pre-gestational metabolic disruption.
- A biphasic model of vascular adaptation to maternal obesity is proposed.

**Graphical abstract:** 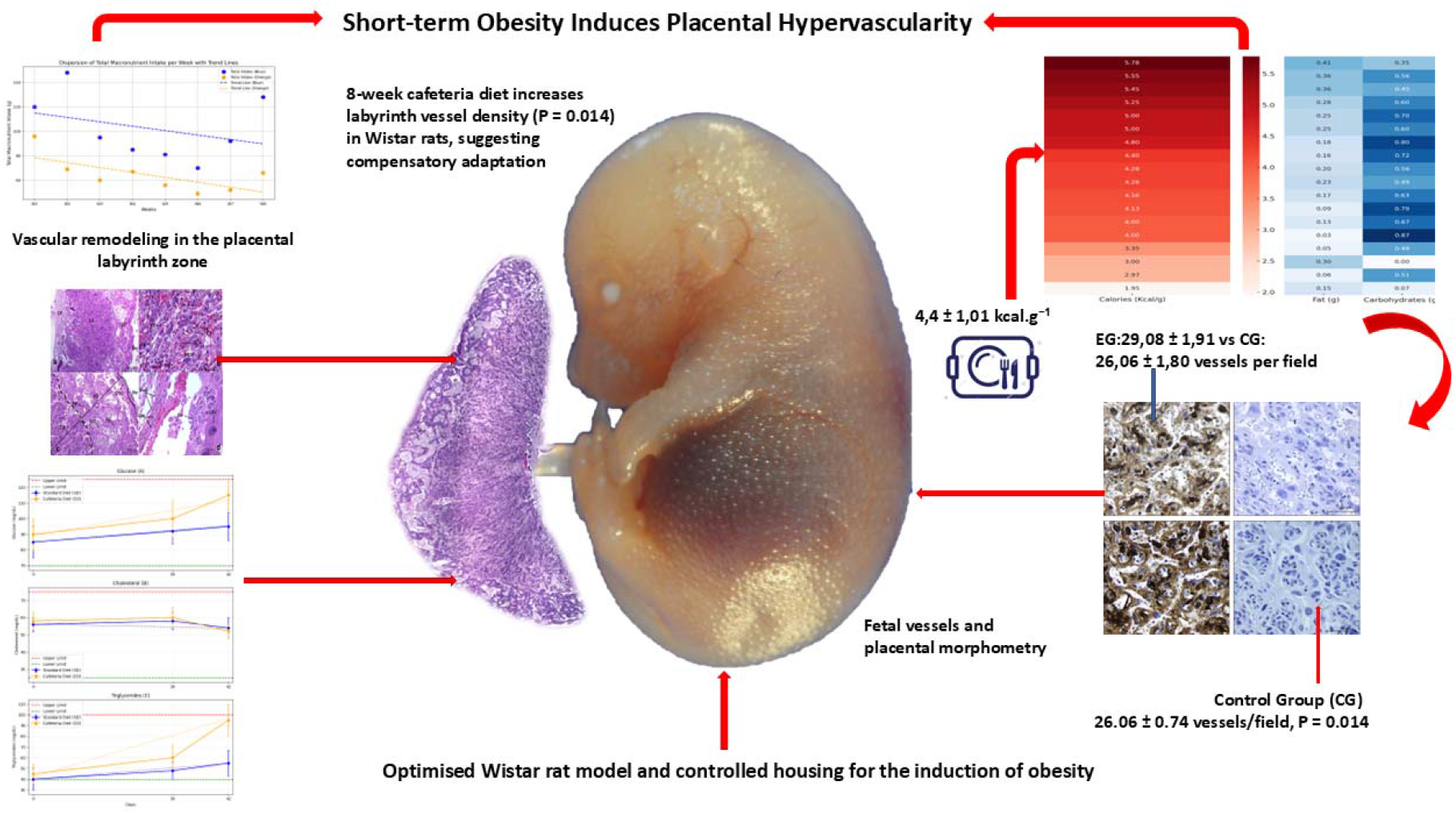

## 1. Introduction

The intricate architecture of the placental vasculature is essential for foetal development, yet it is susceptible to maternal metabolic disturbances [1,2]. Despite the extensive association between chronic maternal obesity and placental hypovascularity, resulting in placental dysfunction, the immediate adaptive responses of the placenta to an acute obesogenic environment remain largely unknown [3]. This knowledge gap is significant because short-term dietary changes can trigger different compensatory mechanisms that precede the well-documented pathologies of long-term obesity [4]. The existence of contradictory findings from various animal models has further complicated our understanding of these initial adaptations, highlighting the need for a more precise model [5,6].

The cafeteria diet (CD) provides a distinctive model for studying the dissociation of rapid adiposity from the systemic metabolic derangements characteristic of chronic obesity models [7,8]. By inducing obesity over a shorter timeframe, this model provides a critical window during which to isolate the placenta’s primary adaptive responses. The hypothesis was formulated that, in contrast to the hypovascularity observed in chronic states, short-term exposure to a hypercaloric, nutrient-imbalanced CD would trigger a compensatory hypervascularization in the placental labyrinth, the primary site of materno-fetal exchange, to ensure the preservation of nutrient transport [9,10].

The aim of this study was to characterise the early vascular remodelling in the placental labyrinth of Wistar rats following an eight-week diet consisting of a high-fat, low-fibre cafeteria diet. The integration of morphometric profiling, biochemical analysis, and targeted CD31/α-actin immunohistochemistry was undertaken with the objective of providing novel insights into the initial plasticity of the placenta in its adaptation to acute maternal metabolic stress.

## 2. Methods

### 2.1 Ethical approval

All experimental procedures involving animals were approved by the Ethics Committee of the Universidad del Valle and conducted in strict accordance with the ARRIVE 2.0 guidelines for animal research [11].

### 2.2 Animal model and diet-induced obesity protocol

Eighteen adult female Wistar rats (11 weeks old; 212.5 ± 22.5 g) were obtained from a certified breeder and acclimatized for a period of one week. The animals were housed individually in a controlled environment (22–23 °C, 12:12-h light/dark cycle) with ad libitum access to water, a configuration that permitted precise monitoring of food consumption. Subsequent to acclimatisation, the rats were randomly assigned to one of two groups (n=9 each) for an eight-week dietary intervention

#### 2.2.1 Control group (CG)

The subjects were provided with a standard rodent diet, which is defined as providing 3.35 kcal/g of energy.

#### 2.2.2 The experimental group (EG)

The subjects were provided with a cafeteria diet that was both hypercaloric and hyperpalatable (CD; 4.4 ± 1.01 kcal/g), consisting of 17 ultra-processed, human-grade food items, which were supplemented with standard diet pellets. This diet was specifically designed to induce rapid obesity through sensory-driven hyperphagia, mirroring human Western dietary patterns [12,13]. The daily food intake and spillage of the subjects were meticulously recorded in order to calculate the energy balance.

### 2.3 Maternal morphometric and metabolic analysis

The development of the obese phenotype was monitored on a weekly basis via morphometric assessments, including body weight (g) and abdominal circumference (cm). At the conclusion of the study (week 8), the presence of abdominal fat accumulation was qualitatively validated in a subset of animals (n=2 per group) using computed axial tomography (CAT) under sedation.

The metabolic status of the subjects was then assessed by the collection of fasting (12-hour) blood samples on days 0, 30, and 44. Glycemia was measured using a glucometer that had been validated for the purpose, while serum cholesterol and triglyceride levels were determined using a commercial colourimetric kit. These were then analysed with a photometer.

### 2.4 Mating, gestation, and placental collection

Subsequent to the 8-week dietary period, female rats were mated with fertile males. On gestational day 0.5, the presence of sperm was confirmed in a vaginal smear, thereby indicating the initiation of gestation. The assigned diets were adhered to throughout the period of gestation. On gestational day 16.5, a time point corresponding to peak placental vascularization, the dams were euthanized via intraperitoneal injection of ketamine (75 mg/kg) and xylazine (10 mg/kg). Feto-placental units were promptly excised, and placental weight, major/minor diameters, and central thickness were documented for macroscopic analysis.

### 2.5 Placental histology and immunohistochemistry (IHC)

The placental tissues were fixed in paraformaldehyde, processed for paraffin embedding, and sectioned at 4 µm. A subset of sections was subjected to staining with Hematoxylin and Eosin (H&E) in order to facilitate evaluation of the general histological architecture of the labyrinth, junctional, and decidual zones.

For the purpose of vascular analysis, an immunohistochemistry (IHC) procedure was conducted in accordance with a dual-staining protocol, the objective of which was to identify both endothelial cells and perivascular support cells. Foetal vessels were labelled with a primary antibody against CD31/PECAM-1 (1:10; MyBiosource), while pericytes and smooth muscle cells were identified with an antibody against α-actin (1:10; Santa Cruz). The slides were imaged at 40X magnification using a Leica DM750 microscope.

### 2.6 Quantification of placental vascularity

Vessel quantification was performed exclusively within the labyrinth zone, the primary site of maternal-foetal exchange. For each placenta, five random, non-overlapping fields were captured. The quantification of foetal vessel density was conducted utilising ImageJ software (NIH) within a standardised area of 1,150,313.4 µm^2^, with the objective of ensuring consistency and comparability across all samples.

### 2.7 Statistical analysis

The analysis of quantitative data was conducted utilising SPSS 22.0 software (IBM), with statistical significance being established at a threshold of P < 0.05. The normality of data distribution was confirmed with the use of Shapiro-Wilk tests, thus validating the application of parametric analyses. Comparisons between the control and experimental groups were conducted using independent samples t-tests, while longitudinal data, such as weekly morphometric changes, were assessed with repeated-measures ANOVA. The results obtained are presented as the mean value ± standard deviation (SD). The complete dataset that was both generated and analysed for the purposes of this study is now available to the public in the Mendeley Data repository. This action was taken in order to ensure transparency and to facilitate the reuse of the data.

## 3. Results

### 3.1. Cafeteria diet drives hyperphagia and macronutrient imbalance

The cafeteria diet (CD), which comprised 17 ultra-processed items, exhibited a distinctly divergent nutritional profile in comparison to the standard diet (SD). The CD was characterised by a higher caloric density (mean 4.4 kcal/g vs. 3.35 kcal/g), elevated fat content (0.21 g/g vs. 0.05 g/g), and a significant increase in carbohydrates (0.55 g/g vs. 0.48 g/g). Conversely, the CD demonstrated a substantial decrease in protein levels (0.06 g/g vs. 0.24 g/g). This hyperpalatable diet successfully induced voluntary hyperphagia in the experimental group (EG), which consumed significantly more calories per day than the control group (CG) (57.34 ± 3.61 kcal/day vs. 34.04 ± 1.75 kcal/day; P < 0.0001). Consequently, the EG consumed substantially more fat (15.65 ± 0.96 g/week vs. 3.87 ± 0.20 g/week; P < 0.0001) but significantly less protein (7.64 ± 0.77 g/week vs. 17.79 ± 0.91 g/week; P < 0.0001) throughout the 8-week intervention. These trends are illustrated in Figures 1 and 2.

**Figure 1.**
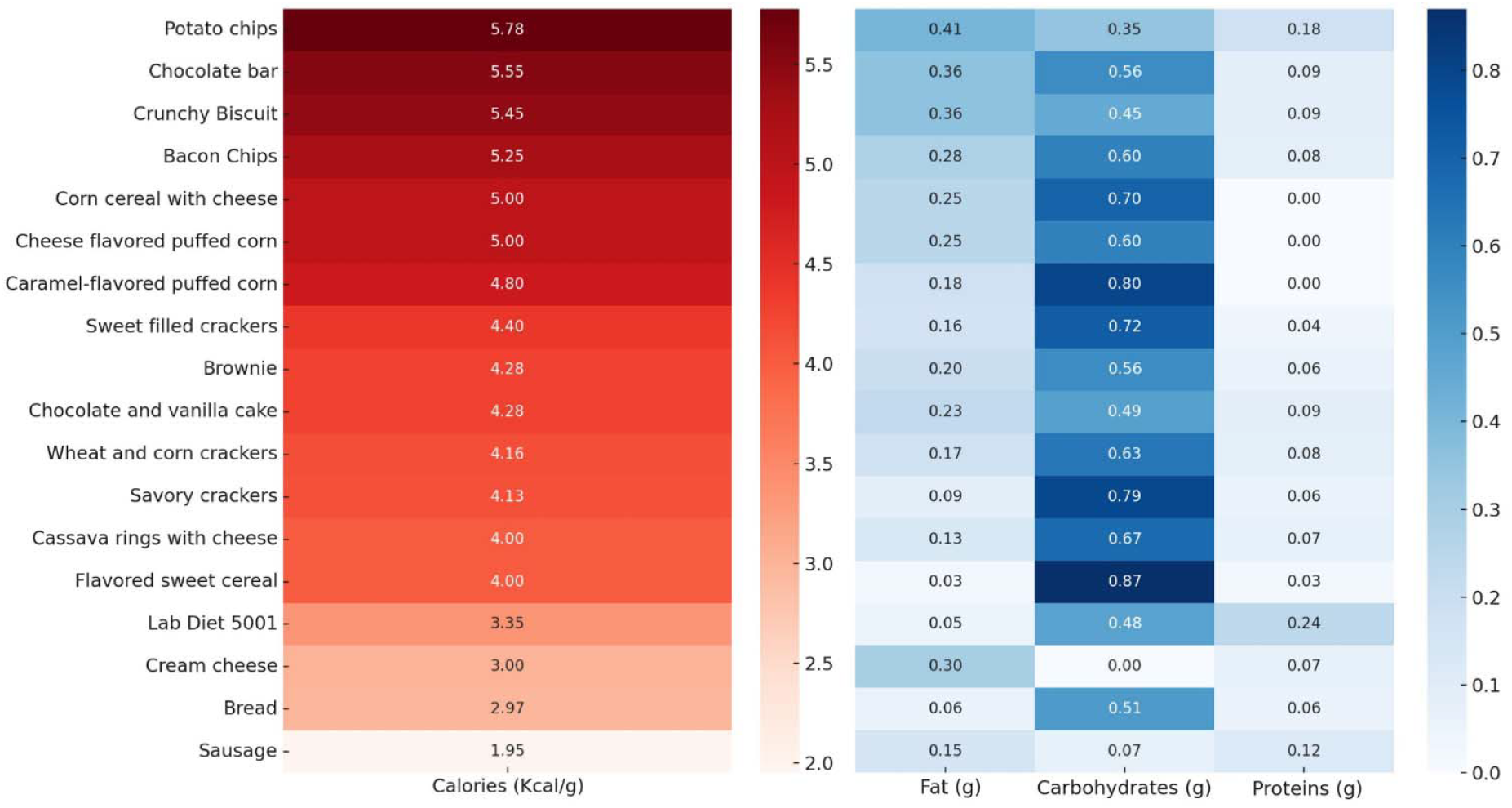
Heatmap comparison of caloric density and macronutrients in cafeteria vs. standard diets. The heatmap contrasts 16 food items from standard (Lab Diet 5001) and cafeteria diets, highlighting caloric density and macronutrient content (fat, carbs, protein in g/g). Red and blue gradients represent kcal/g and macronutrient levels, respectively, enabling rapid visual comparison across diet types.

**Figure 2.**
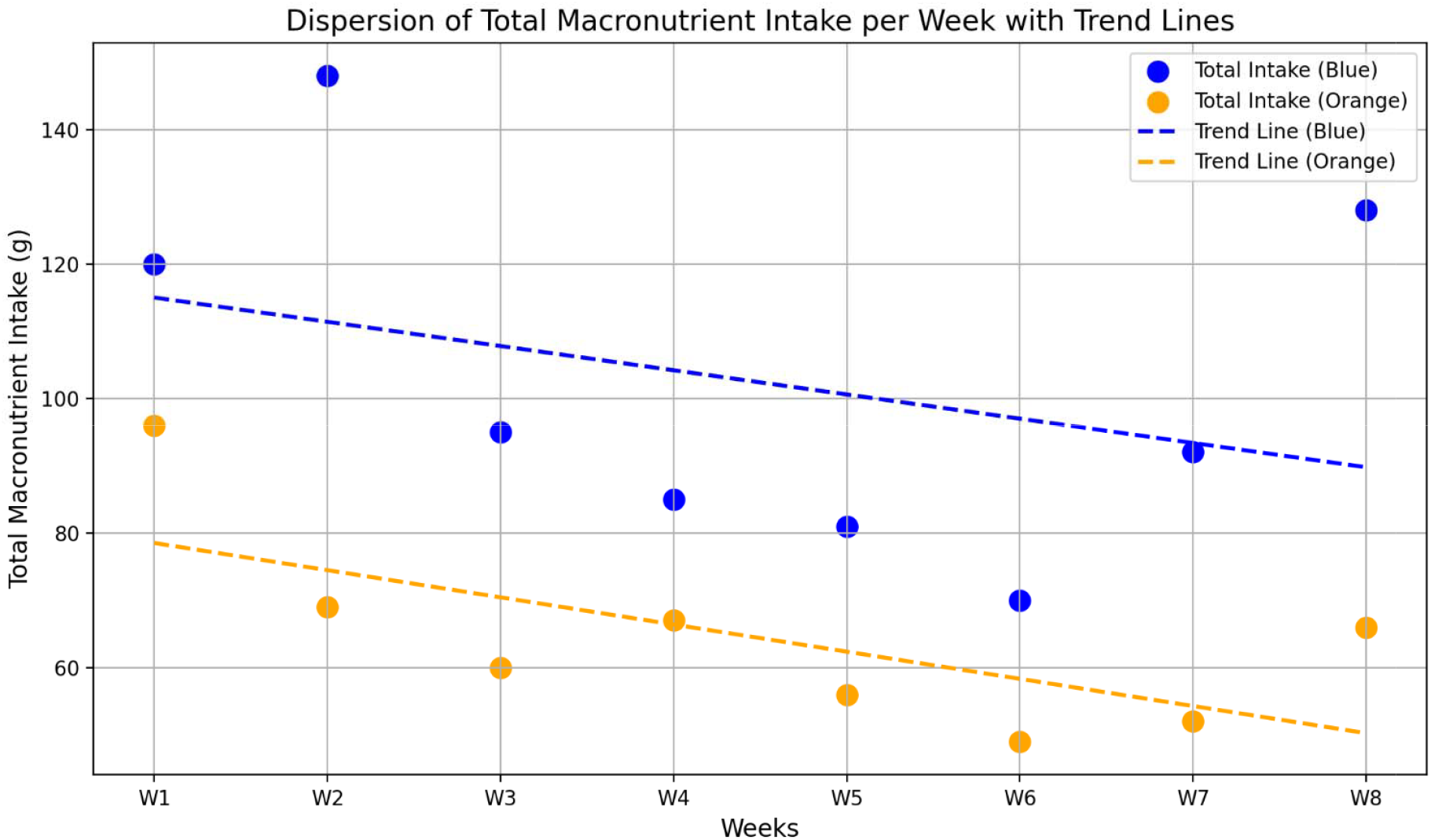
Divergent trajectories of total macronutrient intake over eight weeks with trend lines. The experimental group showed higher overall intake vs. controls (56.69□±□4.34 vs. 45.64□±□2.34□g/week; P□<□0.0001). Intake fluctuated across weeks—elevated in weeks 1–3 and 8, reduced in weeks (P□<□0.0001)—suggesting phase-specific feeding likely driven by satiety or metabolic cues.

### 3.2. A metabolically healthy obese phenotype is induced

The dietary intervention resulted in a significant difference in weight gain. It was observed that, at the commencement of the experiment, the body weights of the rats in the experimental group (EG) and the control group (CG) were comparable (P = 0.31). However, over the course of the eight-week study, the EG rats exhibited a weight gain that was almost double that of the CG rats (62.73 ± 10.93 g vs. 32.26 ± 5.71 g; P < 0.0001). This was accompanied by a significant increase in abdominal circumference, indicative of central adiposity (P < 0.005). As demonstrated by the results of the computed axial tomography scans conducted at week 8, there was a substantial increase in visceral and subcutaneous fat accumulation in the EG. As illustrated in Figure 3, there was no discrepancy in body length, which is an indicator of skeletal growth, between the groups under consideration. Notwithstanding the manifest obese phenotype, no statistically significant disparities were identified in fasting glycemia, cholesterol, or triglyceride levels between the EG and CG at any designated time point (P > 0.05). This finding suggests that the 8-week protocol did not result in systemic metabolic dysfunction (see Figure 4).

**Figure 3.**
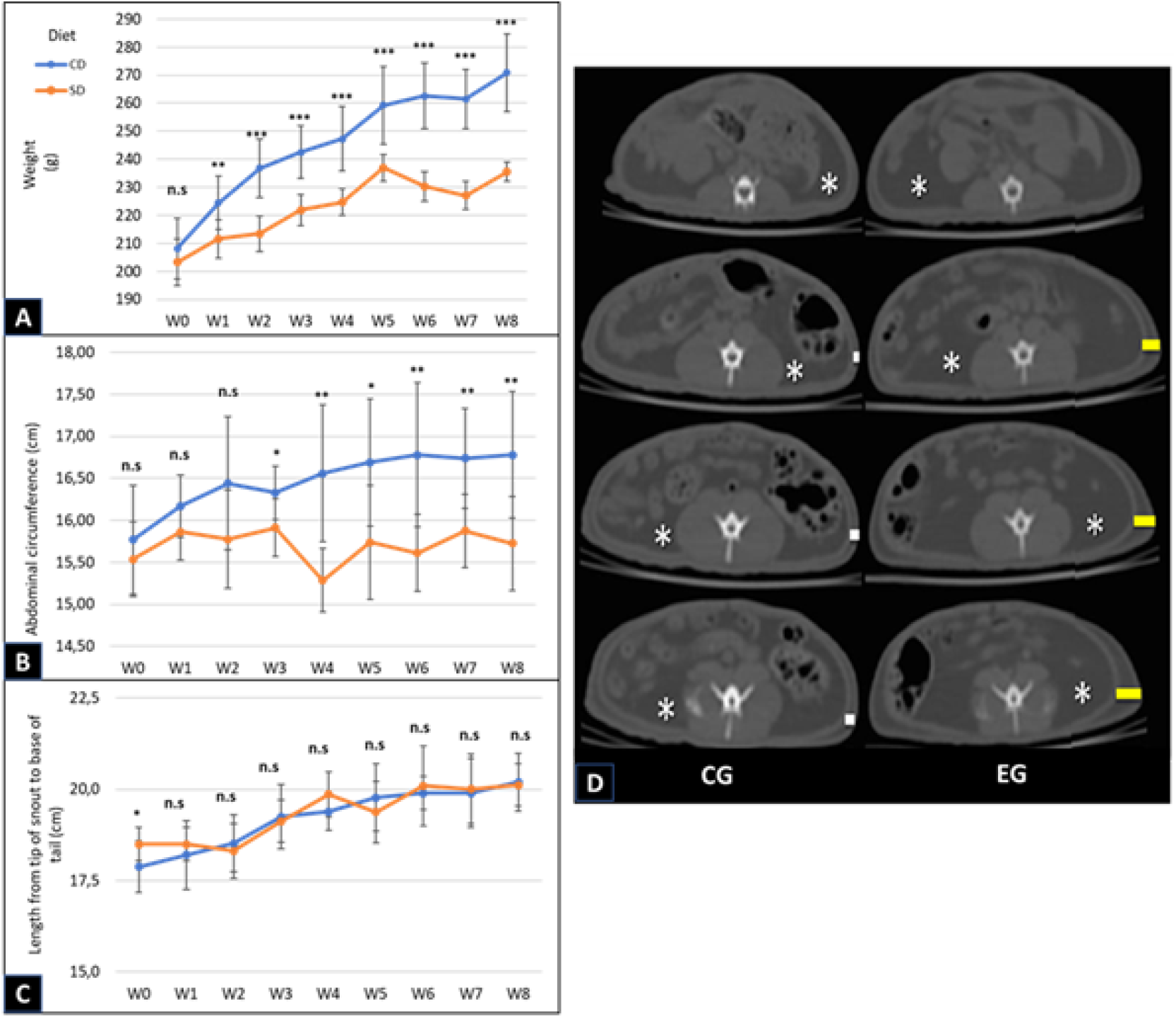
Longitudinal analysis of key morphometric parameters in murine biomodels subjected to either a standard diet (SD, control) or a hyperpalatable cafeteria diet (CD, experimental). Figure 3A–C show weekly trajectories of body weight, abdominal circumference, and body length. CD-fed animals exhibited accelerated weight gain and increased central adiposity (*P*D<D0.0001), with no change in linear growth. Figure 3D presents CT images at endpoint, highlighting visceral (*) and subcutaneous (yellow lines) fat. Data are mean*□*±*□*SD; P values via Student’s t-test (*□<□0.05; **□<□0.005; ***□<□0.0001; n.s.□=□not significant).

**Figure 4.**
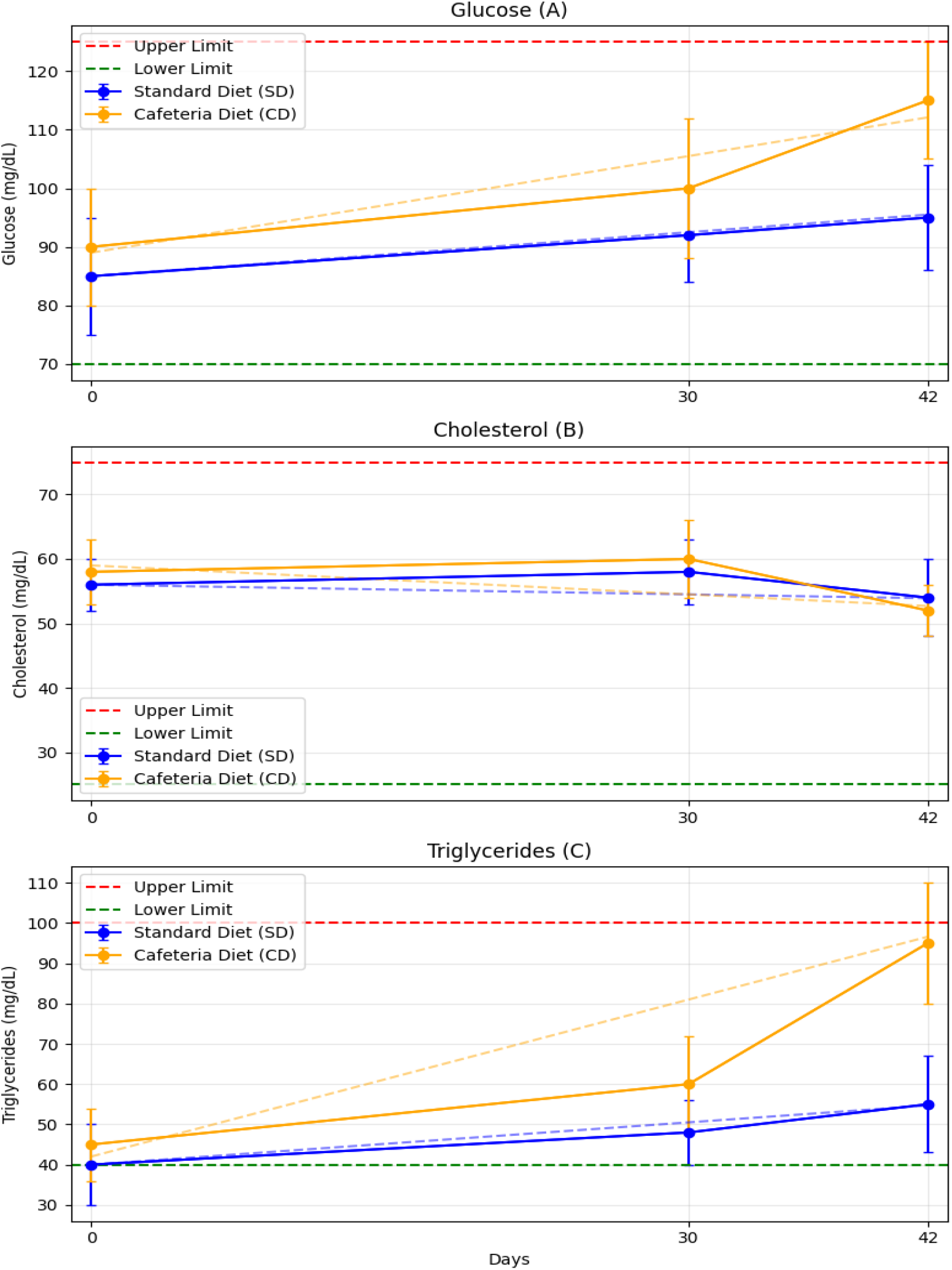
Temporal trends in glucose, cholesterol and triglyceride levels.

### 3.3. Feto-placental growth is preserved despite maternal obesity

At gestational day 16.5, macroscopic examination revealed no significant differences in feto-placental parameters between the groups. The mean foetal weights (EG: 0.41 ± 0.04 g; CG: 0.42 ± 0.07 g; P = 0.60) and placental weights (EG: 0.27 ± 0.05 g; CG: 0.30 ± 0.05 g; P = 0.053) were found to be statistically indistinguishable. Conversely, placental diameters, central thickness, and surface area exhibited no substantial alterations, indicating that short-term maternal obesity did not affect foetal or placental growth at this stage.

### 3.4. Maternal obesity triggers robust hypervascular remodeling in the placental labyrinth

Histological analysis confirmed the preservation of normal placental architecture in both groups, with distinct labyrinth, junctional and decidual zones (see Figure 5). Quantitative histomorphometry revealed no significant differences in labyrinth zone thickness or total placental area between the two groups (P > 0.05).

**Figure 5.**
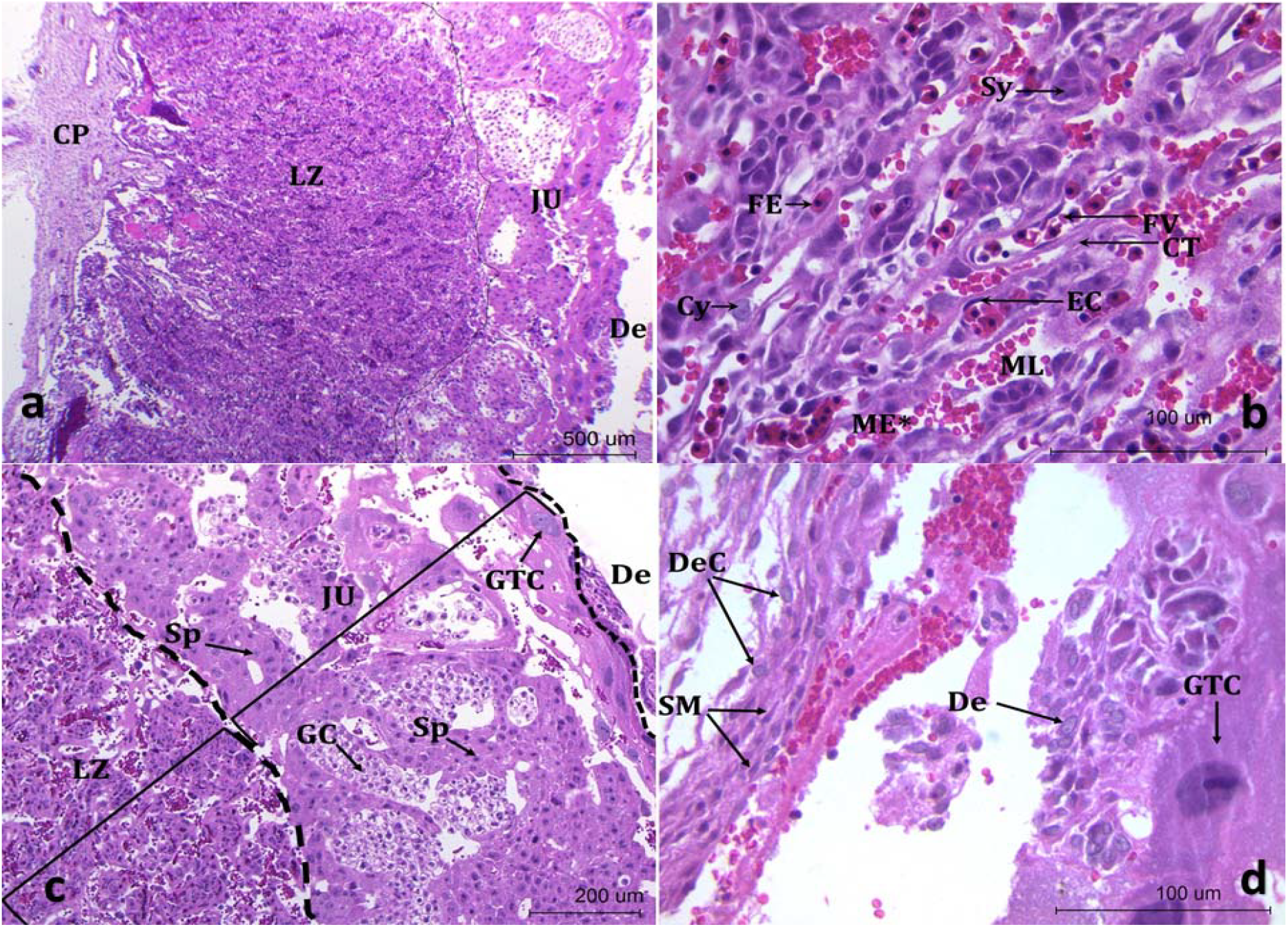
Histological atlas of placental zones at 16.5 dg in control and experimental groups. Figure 5a–d show placental histoarchitecture at increasing magnifications: chorionic plate, labyrinth, junctional zone, and decidua. Key structures include fetal/maternal vessels, trophoblast subtypes, and connective components. Labels: CP, LZ, JU, De, FV, FE, EC, ML, ME*, Sy, Cy, CT, Sp, GC, GTC, DeC, SM.

However, immunohistochemical analysis of the labyrinth – the primary site of maternal-foetal exchange – revealed a significant adaptive change. Furthermore, staining for the endothelial marker CD31 demonstrated a marked increase in foetal vessel density in the EG placentas. Quantitative analysis confirmed this observation, showing that the EG had a significantly higher number of foetal vessels per standardized area compared to the CG (523 ± 33 vs. 469 ± 32 vessels/1,150,313.4 µm^2^; P = 0.014). The finding of labyrinthine hypervascularization, confirmed with negative controls to ensure antibody specificity, is illustrated in Figure 6.

**Figure 6.**
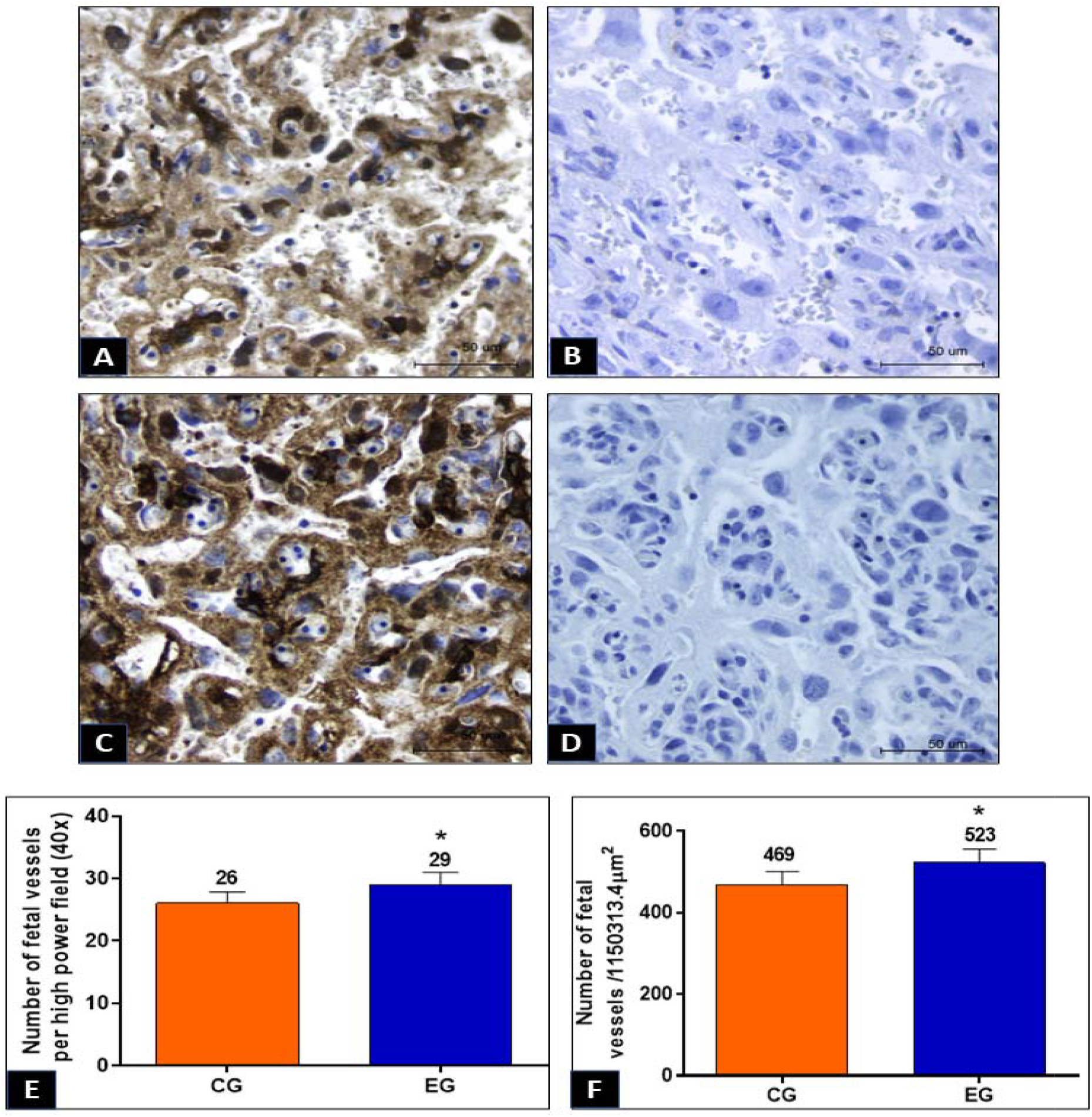
Immunohistochemical profiling of foetal vascular architecture in the labyrinth at 16.5 dg. (A, C) CD31/α-SMA staining in placental labyrinths from control (A) and cafeteria diet (C) groups shows increased vascularity in EG. (B, D) Negative controls confirm antibody specificity. (E, F) Quantification of fetal vessel density per field and area shows significant increase in EG (P□<□0.02; mean□±□SD; Student’s t-test).

## 4. Discussion

The most significant finding of this study is the induction of hypervascularity in the placental labyrinth of Wistar rats after a short-term cafeteria diet. This finding is counterintuitive, as it contradicts the substantial body of literature documenting vascular deficits, endothelial dysfunction and reduced capillary density in placental tissue from pregnancies affected by chronic obesity or type 2 diabetes [14–16]. Such chronic conditions are generally associated with systemic inflammation, oxidative stress and advanced glycation end-products, all of which have been demonstrated to suppress angiogenesis and promote vascular degeneration [17–19]. The present study provides compelling evidence for the existence of a biphasic placental vascular response to obesity-promoting stress, comprising an initial proliferative phase, which may be adaptive, that likely precedes the overt vasculopathy seen in long-term metabolic disease. This distinction highlights the importance of considering the temporal and metabolic context of the maternal stressor when determining the placental phenotype [19,20].

It is proposed that this robust angiogenic response is a direct consequence of the unique physiological milieu created by the experimental model [6,8,21]. The cafeteria diet, which effectively mimics the palatable, energy-dense, and nutritionally imbalanced nature of a human “Western” diet [22,23], induced a rapid and substantial increase in maternal adiposity within a mere eight weeks (Figure 3). It is important to note that this obese phenotype was established without concurrent systemic metabolic derangements, such as fasting hyperglycaemia or dyslipidaemia (see Figure 4). This state of “metabolically healthy” obesity, suggests that key signalling pathways, including insulin sensitivity, may have been largely preserved [24,25]. This may have enabled the placenta to retain its inherent plasticity and mount a powerful compensatory response. Furthermore, the specific composition of the diet, notably its low protein content (Figure 1), is a potent, independent stimulus for this remodelling. Nutrient-sensing pathways, including the mammalian target of rapamycin (mTOR) and general control nonderepressible 2 (GCN2), exhibit heightened sensitivity to amino acid availability [26,27]. A low-protein signal has been shown to trigger these pathways, leading to the upregulation of angiogenic factors such as Vascular Endothelial Growth Factor (VEGF) [17,19]. This process is believed to be a physiological response aimed at increasing the surface area available for transport and maximising the uptake of limiting amino acids, which are essential for foetal development [1,5,28].

A critical component of our findings is the evident dissociation between the marked increase in foetal vessel density (see Figure 6) and the absence of any corresponding increase in foetal or placental weight. This finding indicates that the observed relationship between vessel quantity and enhanced vascular function may not be direct. The process of rapid, demand-driven angiogenesis, has the potential to result in a structurally compromised vascular network [29,30]. For instance, while an increase in CD31-positive endothelial cells was observed, the maturation and stability of these new vessels is contingent on adequate recruitment of perivascular cells, such as pericytes (stained here by α-actin). It is conceivable that the proliferation of endothelial tubes may exceed the migration and investment of these supporting cells, resulting in a network of capillaries that are immature, leaky, and hemodynamically inefficient [18]. Such a disorganized capillary bed, despite its density, would fail to improve and could even impair the efficiency of maternal-foetal exchange, providing a mechanistic explanation for the lack of accelerated foetal growth [5,31].

The consequences of this early placental maladaptation for developmental programming are significant. It is evident that an immature and inefficient vascular architecture [17], even if only transient, has the capacity to effect a fundamental alteration to the in utero environment. This may result in subtle but chronic foetal hypoxia or an imbalanced flux of nutrients, which can lead to programming of the foetal metabolic and cardiovascular systems, thereby increasing the risk of disease in later life [9,32]. This early adaptive phase, therefore, represents a critical and hitherto unappreciated window for intervention. Non-pharmacological strategies, such as maternal exercise [33], have been demonstrated to enhance nitric oxide bioavailability and promote a shift from disorganised angiogenesis to mature, stable vascular networks (arteriogenesis). From a pharmacological standpoint, targeting the metabolic dysfunction is also pivotal [2,8]. The high saturated fat content of the cafeteria diet is likely to induce placental lipotoxicity. Interventions with peroxisome proliferator-activated receptor gamma (PPARγ) agonists, have shown promise in restoring placental fatty acid oxidation and normalising angiogenic signalling in obesogenic states [29,30]. It is also imperative for future studies to stratify analyses by foetal sex [34,35], as the placenta’s response to metabolic insults is known to be highly dimorphic, a critical factor that may have been masked in our pooled analysis.

In conclusion, the present study provides compelling evidence that short-term, diet-induced maternal obesity triggers a compensatory increase in vascularity in the rat placental labyrinth. This early adaptive response occurs prior to systemic metabolic collapse and does not enhance foetal growth, suggesting a potential decoupling of vascular structure and function. The present study proposes a biphasic model of placental adaptation to maternal obesity, beginning with this angiogenic phase. It is imperative to delineate the mechanisms, functional consequences, and long-term implications of this initial vascular remodelling in order to formulate strategies to mitigate the intergenerational cycle of metabolic disease.

